# CRISPR Turbo Accelerated Knock Out (CRISPy TAKO) for rapid *in vivo* screening of gene function

**DOI:** 10.1101/2020.08.04.236968

**Authors:** SL Plasil, A Seth, GE Homanics

## Abstract

The development of CRISPR/Cas9 technology has vastly sped up the process of genome editing by introducing a bacterial system that can be exploited for reverse genetics-based research. However, generating homozygous knockout (KO) animals using traditional CRISPR/Cas9-mediated techniques requires three generations of animals. A founder animal with a desired mutation is crossed to produce heterozygous F1 offspring which are subsequently interbred to generate homozygous F2 KO animals. This study describes a novel adaptation of the CRISPR/Cas9-mediated method to develop a homozygous gene-targeted KO animal cohort in one generation. A well-characterized ethanol-responsive gene, *MyD88*, was chosen as a candidate gene for generation of *MyD88*^*-/-*^ mice as proof of concept. Previous studies have reported changes in ethanol-related behavioral outcomes in MyD88 KO mice. Therefore, it was hypothesized that a successful one-generation KO of MyD88 should reproduce decreased responses to ethanols sedative effects, as well as increased ethanol consumption in males that were observed in previous studies. One-cell mouse embryos were simultaneously electroporated with four gRNAs targeting a critical Exon of MyD88 along with Cas9. DNA and RNA analysis of founder mice revealed a complex mix of genetic alterations, all of which were predicted to ablate MyD88 gene function. This study additionally compared responses of Mock treatment control mice generated through electroporation to controls purchased from a vendor. No substantial behavioral changes were noted between control cohorts. Overall, the CRISPR/Cas9 KO protocol reported here, which we call <u>CRISP</u>R <u>T</u>urbo <u>A</u>ccelerated <u>K</u>nock<u>O</u>ut (CRISPy TAKO), will be useful for reverse genetic *in vivo* screens of gene function in whole animals.

## Introduction

Clustered regulatory interspaced short palindromic repeats (CRISPR) paired with CRISPR associated protein 9 (Cas9) is currently the dominant and preferred gene editing tool in scientific research. CRISPR based screens of gene function *ex vivo* have been tremendously useful for identifying genes involved in tumor suppression^1^, mitochondrial function^2^, and dendritic development^3^. High throughput CRISPR loss-of-function reverse genetic screens allow for the rapid identification of genes involved in phenotypes of interest. However, *in vitro* screens are limited by phenotypes that can be readily assayed in cell culture, e.g., cellular proliferation, drug sensitivity, and cell survival. Further, the acquisition of transcriptome data has greatly outpaced our capacity to functionally study genes of interest. For many biological questions, particularly those that pertain to dysfunction of the central nervous system where behavioral abnormalities are the primary phenotype of interest, *in vivo* tests of behavior must be employed. Because behavior is the phenotype of interest, *in vitro* screens are unsatisfactory. In this study, we sought to develop a method with moderately high throughput that could be used *in vivo* to screen genes for effects on behavior.

Global gene knockout (KO) animal models are a gold standard approach that have been widely used to study and delineate the effects of individual molecules in whole organisms. The recent application and widespread adoption of CRISPR/Cas9 technology dramatically facilitated KO animal generation. However, the standard method of creating CRISPR KO animals, a.k.a., CRISPy Critters^4^, typically requires three generations to produce experimental animals that can be phenotypically evaluated and therefore is unsuitable for moderate-high throughput *in vivo* screens (Fig. 1A). Briefly, in a typical CRISPR KO animal study, CRISPR reagents are introduced to one-cell embryos that develop into founder (F0) animals that are screened for the desired mutation. F0 animals are typically an eclectic mix of wild-type and mutant animals. The mutants may be heterozygotes, homozygotes, or compound heterozygotes, and most mutant alleles differ in the individual mutations they harbor in the target gene of interest. A founder animal that harbors a desirable mutation (typically a frameshift or a large deletion) is then mated to wild-type (WT) mice to produce heterozygous F1 offspring. Subsequently, heterozygotes are interbred to produce homozygous F2 mutant KO offspring. These F2 mutant animals have both alleles of the gene of interest inactivated, they all harbor the same mutation in the gene of interest, and they can be compared to WT littermate controls for relevant phenotypic changes. Although this CRISPR approach to creating gene KO animals represents a dramatic savings in time, effort, and expense compared to traditional embryonic stem cell based gene targeting approaches, the CRISPR approach still requires considerable time and expense because three generations of animal production is time consuming and results in substantial animal care and housing expenses. This process also requires a considerable amount of personnel time for colony maintainance and genotyping.

**Figure 1:**
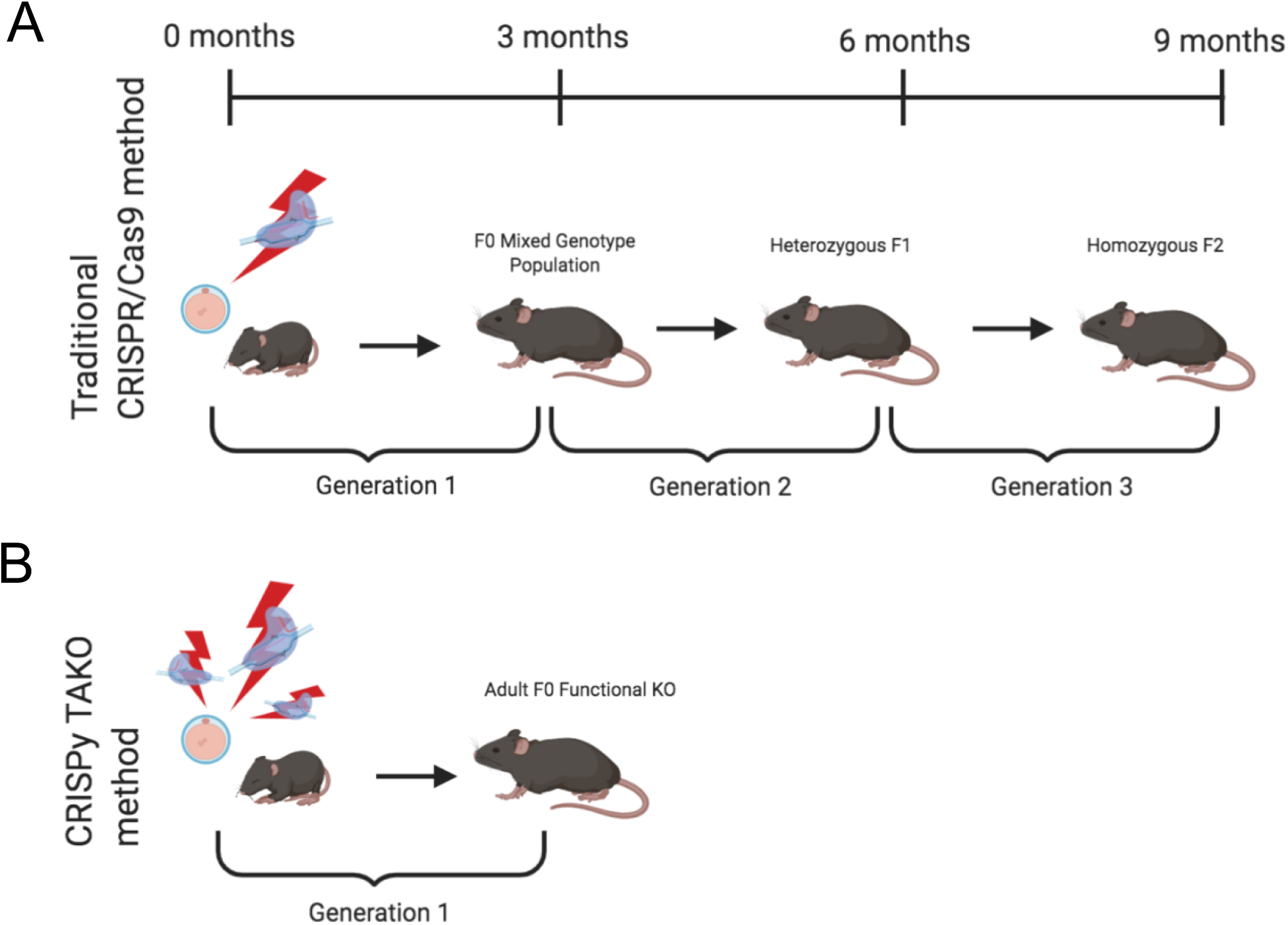
Comparison of the timeline required to produce knockout (KO) mice for behavioral testing using traditional and the CRISPy TAKO approaches. **(A)** Traditional CRISPR/Cas9-mediated method to create a stable KO line. Founder (F0) animals are an eclectic mix of wild-type, heterozygous, and homozygous KOs. A founder with an inactivating mutation is selected for breeding to establish a KO line of mice. First generation offspring (F1) are heterozygous and must be interbred to produce an F2 generation. A subset (∼25%) of the F2 generation are homozygous KO mice and can be compared for behavioral phenotypes with WT littermates. **(B)** CRISPy TAKO method for creating functional KO mice. By using multiple gRNAs that target a small but functionally critical part of the gene of interest, most F0 mice harbor biallelic mutations that functionally inactivate the gene of interest and are suitable for behavioral phenotyping.

We endeavored to establish a one generation CRISPR KO approach in which F0 animals could be directly used to test for the behavioral consequences of gene inactivation. We reasoned that a very high efficiency CRISPR mutagenesis approach could be used to efficiently create F0 animals in which both alleles of the gene of interest are mutated and are functionally inactivated (i.e., gene KOs) (Fig. 1B). Although each F0 animal may have different mutations, they would all be functionally and phenotypically equivalent if a critical part of the gene were sufficiently mutated.

Our long-term goal is to employ this accelerated technique to vastly speed up the screening process of testing novel ethanol-responsive genes for involvement in ethanol-related behavioral phenotypes, including ethanol consumption. Therefore, we initially piloted this approach *in vitro* on two novel ethanol-responsive long noncoding RNA (lncRNA) genes. We subsequently sought to validate this method *in vivo* by mutating a gene previously shown to alter behavioral responses to ethanol when inactivated using traditional global KO technology. *MyD88* was chosen as a well-characterized ethanol-responsive gene for proof-of-concept as prior studies have evaluated the effects of MyD88 global KO on ethanol-related behaviors, including ethanol drinking^5^ and response to ethanol’s acute sedative/hypnotic and motor ataxic effects^6,7^. Single generation F0 MyD88 KO animals were hypothesized to exhibit decreased ethanol-induced sedative/hypnotic effects, decreased sensitivity to ethanol-induced motor ataxia, and a male-specific increase in ethanol consumption relative to controls.

To further streamline this accelerated KO mouse protocol, we reasoned that for first pass screening of genes for behavioral phenotypes, isogenic animals purchased directly from a vendor could be used as a control group for comparison to KOs. However, one concern is that the CRISPR procedure itself, irrespective of the gene being mutated, could exert deleterious effects that could lead to false positive or negative results. Therefore, we also created in-house Mock treatment controls that were produced under an identical protocol to the KOs except that the Mock-treated animals were created with procedures that lacked crRNAs. This Mock-treated control group was directly compared to isogenic C57BL/6J WT mice (Jax controls) purchased from the Jackson Laboratory (JAX). We hypothesized that these two control groups would not differ on behavioral endpoints of interest.

In this report, we describe implementation and validation of a novel technique for the accelerated production of CRISPR KO mice in one generation. Animals produced via this protocol are herein affectionately referred to as <u>CRISP</u>R <u>T</u>urbo <u>A</u>ccelerated <u>K</u>nock<u>O</u>uts (i.e., CRISPy TAKOs). We report that our CRISPR protocol can reliably produce a large number of F0 KO animals and that the ethanol phenotype of MyD88 CRISPy TAKOs largely recapitulates results previously reported for traditional MyD88 global KOs. Furthermore, for the behaviors tested in this study, vendor purchased mice and Mock treatment controls did not differ substantially. Together, these results establish the CRISPy TAKO method for screening gene function *in vivo*. This method has moderately high throughput and will be especially useful for phenotypes, such as behavioral responses, that cannot be assayed *in vitro*.

## Materials and Methods

### Animals

All experiments were approved by the Institutional Animal Care and Use Committee of the University of Pittsburgh and conducted in accordance with the National Institutes of Health Guidelines for the Care and Use of Laboratory Animals. C57BL/6J male and female mice used to generate embryos for electroporation and the purchased control group were procured from The Jackson Laboratory (Bar Harbor, ME). CD-1 recipient females and vasectomized males were procured from Charles River Laboratories, Inc. (Wilmington, MA). Mice were housed under 12-hr light/dark cycles, with lights on at 7 AM and had *ad libitum* access to food (irradiated 5P76 ProLab IsoProRMH3000; LabDiet, St. Louis, MO) and water.

### gRNA Design

Guide RNAs (gRNAs) were generated using a commercially available two-piece system termed ALT-R™ CRISPR/Cas9 Genome Editing System (IDT DNA, Coralville, IA). This system combines a custom CRISPR RNA (crRNA) for genomic specificity with an invariant trans-activating crRNA (tracrRNA) to produce gRNAs^4^. crRNAs were designed using the computational program CCTop/CRISPRator^8,9^, which gauges candidate sgRNAs for efficiency and specificity.

Four crRNAs were used to target the ethanol-responsive lncRNA gene *4930425L21Rik* (see Table 1 for gRNA target sequences). These four crRNAs bind within a 366bp region that includes the putative promoter and first Exon (see Fig. 2A). Similarly, four crRNAs were used to target the lncRNA gene *Gm41261* (see Table 1 for gRNA target sequences). These four crRNAs bind with a 316bp region that includes the putative promoter and first Exon (see Fig. 2C). Four crRNAs were also selected for *MyD88* (see Table 1 for gRNA target sequences) that bind within a 209bp region that includes *MyD88* Exon 3 and flanking DNA (see Fig. 3A). For each project, the four crRNAs were annealed separately with tracrRNA in a 1:2 molar ratio then combined into a single solution.

**Table 1:** Comparison of TAKO cohort results with previous findings.

|  | CRISPy TAKO |  | Previous Reports |  |
| --- | --- | --- | --- | --- |
|  | Males | Females | Males | Females |
| Rotarod | No change | Faster recovery | Faster recovery<sup>6,22</sup> | Faster recovery<sup>6</sup> |
| EtOH Loss of Righting Reflex | Reduced duration | No change | Reduced duration<sup>6,22</sup> | Reduced duration<sup>6</sup> |
| 2BC-EOD (15% v/v) | Increased ethanol drinking | Reduced total fluid intake | Increased ethanol drinking and preference.<br>Reduced total fluid intakes | Reduced total fluid intakes<sup>5</sup> |

**Figure 2.**
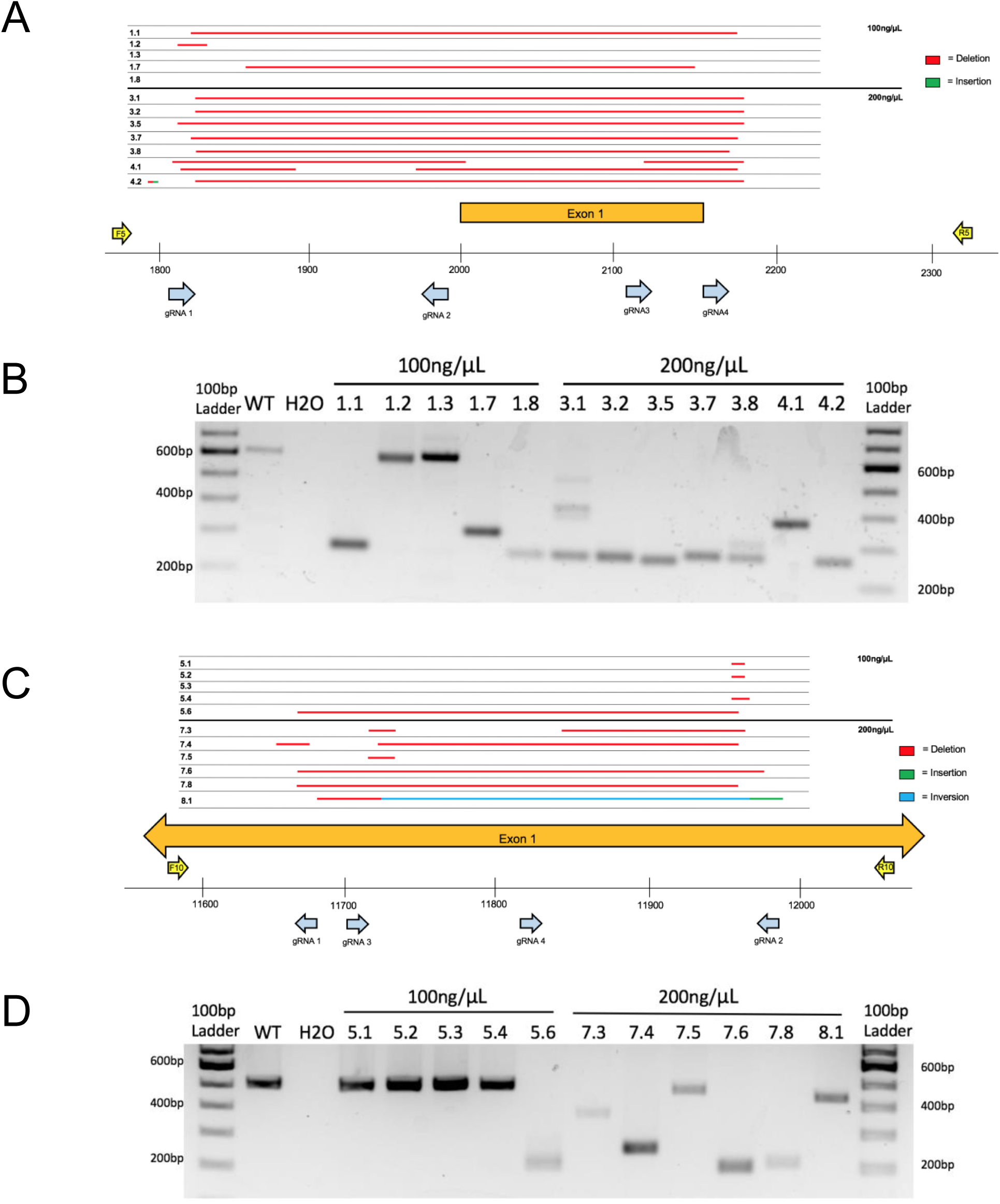
Embryo CRISPy TAKO genotypes for *4930425L21Rik* and *Gm41261*. (A) Sequence results for the major product(s) of TAKO embryos targeting gene *4930425L21Rik* electroporated with 100ng/µL and 200ng/µL Cas9 protein. Full deletions are shown in red. Sequence insertions are shown in green. Individual animal tag numbers are presented on the left. The gRNAs and PCR primers used are shown as blue and yellow arrows, respectively. (B) Agarose gel electrophoresis of PCR amplicons for *4930425L21Rik* in embryos. Samples 1.1 through 1.8 were electroporated with 100ng/µL Cas9 protein while samples 3.1 through 4.2 were electroporated with 200ng/µL Cas9 protein. (C) Sequence results from TAKO embryos targeting gene *Gm41261* with 100ng/µL and 200ng/µL Cas9. Sequence inversions are shown in blue. (D) Agarose gel electrophoresis of PCR amplicons for *Gm41261* in embryos. Samples 5.1 through 5.6 were embryos electroporated with 100ng/µL Cas9 while samples 7.3 through 8.1 were electroporated with 200ng/µL Cas9 protein.

**Figure 3:**
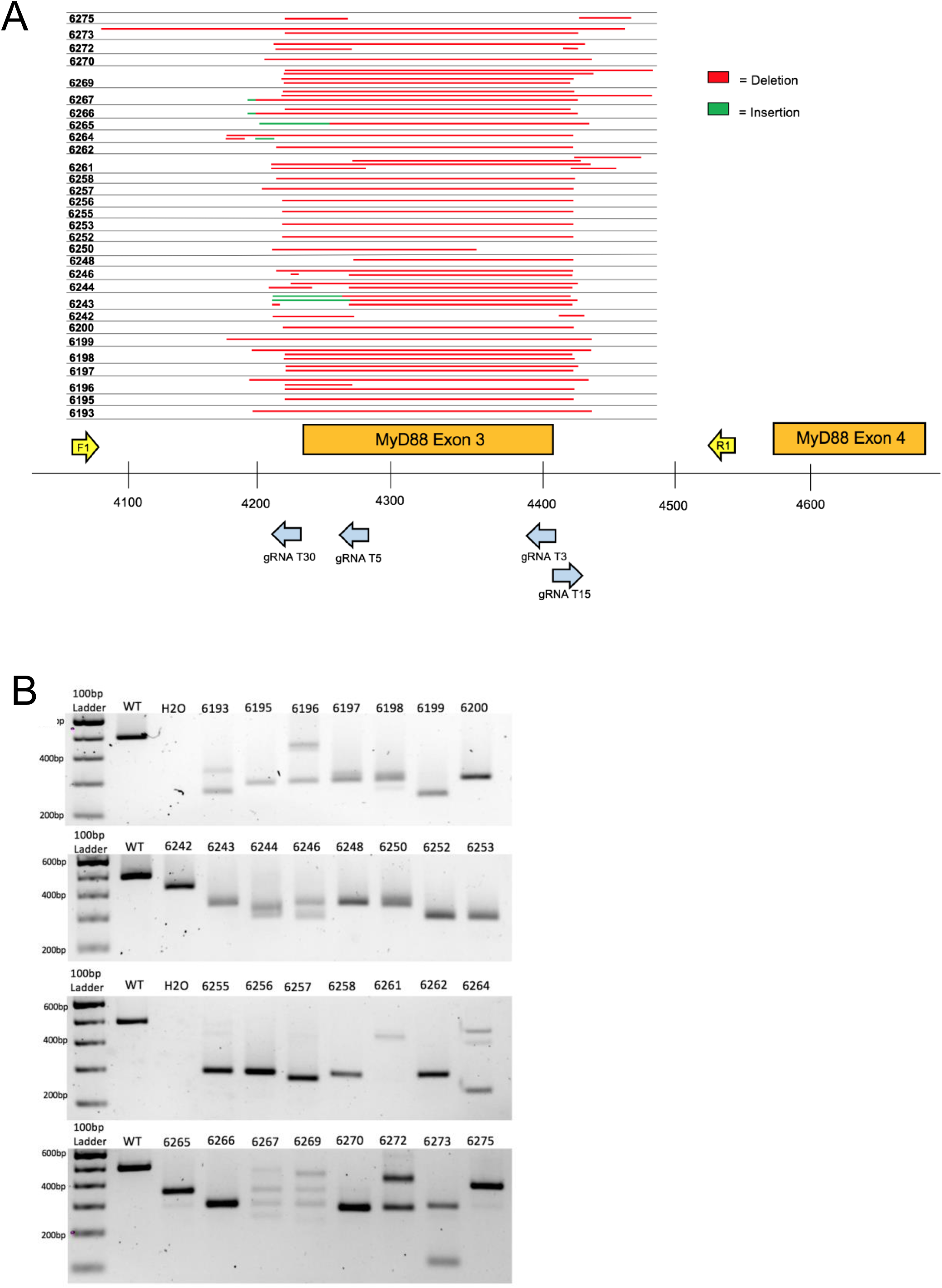
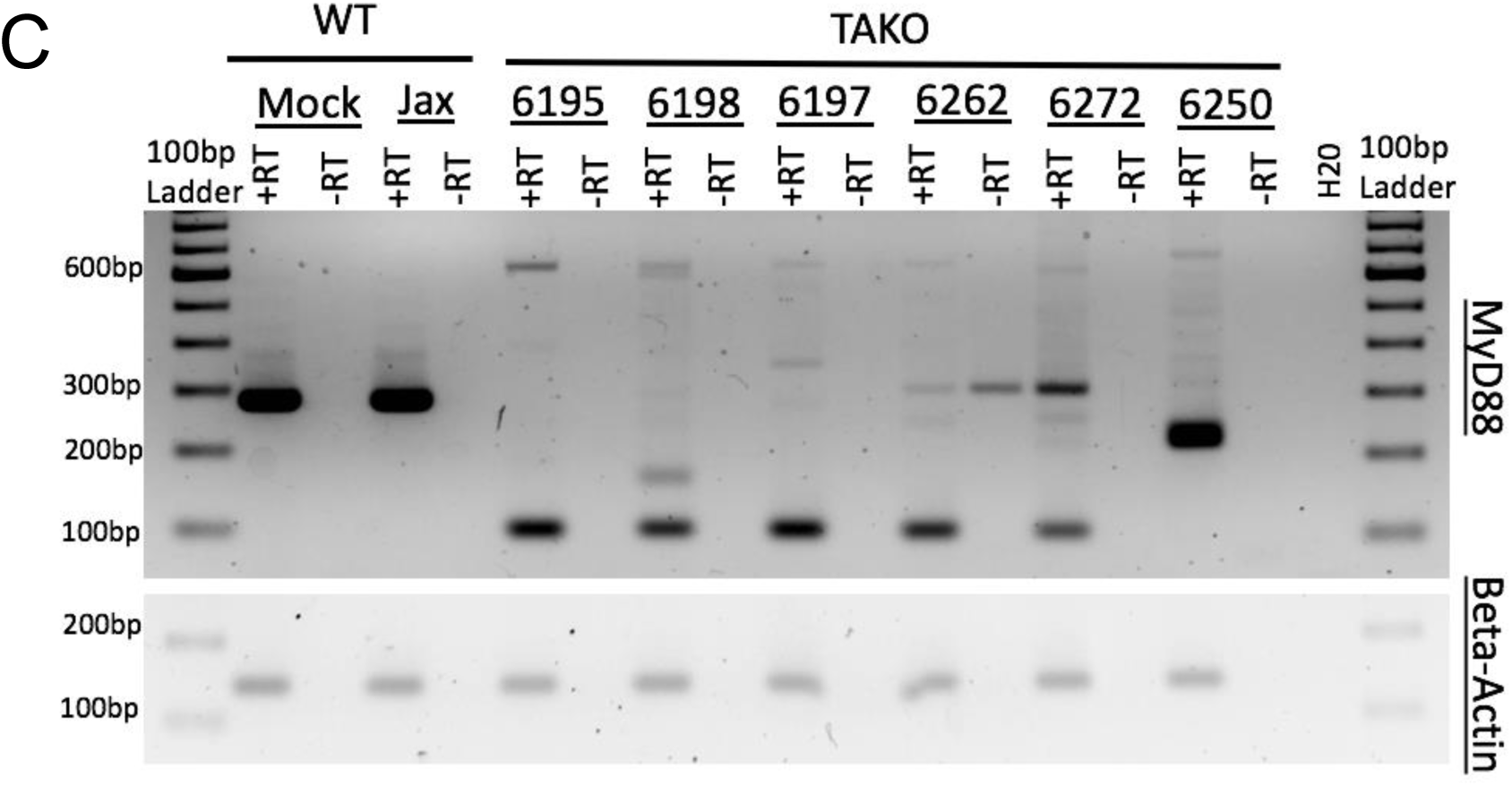
*In vivo* MyD88 CRISPy TAKO genotypes. (A) Sequencing results from mice selected for behavioral experimentation. CRISPR/Cas9 mutagenesis of MyD88 exon 3 was performed and animals used for experimentation were sequenced to confirm successful deletion. Full deletions are shown in red. Sequence insertions are shown in green. Individual animal tag numbers are presented on the left. The gRNAs and PCR primers used are shown as blue and yellow arrows, respectively. (B) PCR results from DNA of MyD88 CRISPy TAKO mice used for experimentation. Jax used as WT control. (C) RT-PCR results from cerebellar brain tissue from a random subset of Mock, Jax, and MyD88 CRISPy TAKO mice showing abnormal *MyD88* RNA transcripts in TAKO mice. RT-PCR using Beta-actin was used as a control.

### CRISPR/Cas9 Mutagenesis

Female C57BL/6J mice were superovulated with 0.1mL of CARD HyperOva (CosmoBio, #KYD-010) between 10 AM and 11 AM, followed by 100 IU of human chorionic gonadotropin (Sigma, #CG10) 46-48hrs later. Donor females were caged overnight with C57BL/6J males starting 4-6hrs post-gonadotropin injection and allowed to mate. Embryos were harvested from oviducts between 9 AM and 10 AM the following morning, cumulus cells were removed using hyaluronidase, and embryos were cultured under 5% CO^2^ in KSOM medium (Cytospring, #K0101) for 1-2hrs. Embryos were electroporated in 5µL total volume of Opti-MEM medium (ThermoFisher, #31985088) containing 100ng/µL of each sgRNA and 100 or 200ng/µL Alt-R® S.p. Cas9 Nuclease V3 protein (IDT, #1081058) with a Bio-Rad Gene-Pulser Xcell in a 1mm-gap slide electrode (Protech International, #501P1-10) using square-wave pulses (five repeats of 3msec 25V pulses with 100msec interpulse intervals). Two different concentrations of Cas9 protein were used to assess which produced greater mutagenesis in embryos targeting genes *4930425L21Rik* and *Gm41261*; only the 200ng/µL concentration was used for MyD88 embryos. Electroporated embryos were placed back into culture under 5% CO^2^ in KSOM. For *4930425L21Rik* and *Gm41261*, embryos were cultured for 3 days until the morulea/blastocyst stage and subsequently analyzed for mutations. For MyD88, one-or two-cell embryos were implanted into the oviducts of plug-positive CD-1 recipient (20-40 embryos per recipient) that had been mated to a vasectomized male the previous night. Mock-treated controls were manipulated in parallel as described above, except that the electroporation mix lacked the *MyD88*-specific crRNAs (i.e., only tracrRNA and Cas9 protein were used).

### Genotyping

For *4930425L21Rik* and *Gm41261*, DNA was amplified from individual embryos using a Qiagen Repli-G kit (Qiagen, #150025) and subject to PCR genotyping under the following settings: 95°C for 5min (1x); 95°C for 30sec, 60°C for 30sec, 72°C for 1min (40x); 72°C for 10min (1x). Primers for PCR amplification of *4930425L21Rik* and *Gm41261* are listed in Table 1. PCR amplicons (WT = 613 anf 506bp, respectively) were analyzed by agarose gel electrophoresis and Sanger sequencing.

For *MyD88*, DNA was isolated from tail snips of MyD88 CRISPy TAKO and Mock-treated control offspring using Quick Extract (Lucigen, #QE09050). Primers for *MyD88* genotyping are listed in Table 1. PCR amplicons (WT = 494bp) were analyzed by agarose gel electrophoresis and Sanger sequencing.

### Subcloning

Samples that did not produce clear chromatograms were subcloned to identify allelic variants. The TOPO™ TA cloning kit (ThermoFisher Scientific, #K457501) was used according to manufacturers instructions, with slight modifications. Briefly, sample PCR product was incubated at room temperature for 15mins with TOPO reagents, then the TOPO vector mixture was incubated with chemically competent DH5α (ThermoFisher Scientific, #18265017) cells on ice for 30min. Cells were then heat-shocked for 45sec in a 42°C water bath then immediately placed back on ice for 2min. S.O.C. medium (ThermoFisher Scientific, #15544034) was added and cells were incubated in a bacterial shaker at 37°C for 90min at 225rpm. Cells were plated on kanamycin-resistent LB plates and incubated at 37°C for 16-18hrs. Single colonies (n = 10 per sample) were collected and their DNA was used for PCR. Colonies that produced a single PCR band were then Sanger sequenced.

### RNA Preparation

Brain cerebellar tissue from one Mock treatment control (n = 1 male), one Jax control (n = 1 male), and 6 MyD88 mutants (n = 3 male, n = 3 female) were used for RT-PCR analysis. All mice were 11-12 weeks of age at time of sacrifice. Total RNA was isolated using TRIzol (Invitrogen, #15596018) according to the manufacturer’s protocol, and purified with a TURBO DNA-*free*™ Kit (Invitrogen, #AM1907). Total RNA was analyzed for purity and concentration using a Nanodrop Spectrophotometer (Thermo Scientific, Waltham, MA). One microgram of purified RNA was converted into cDNA using Superscript™ III First-Strand Synthesis System (Invitrogen, #18080051) with random hexamer primers. PCR primers were used that span from Exon 2 to Exon 4 (see Supplementary Table 1) of MyD88. A reaction that lacked reverse transcriptase was used as a negative control for each sample tested. RT-PCR amplicon size is 280bp for WT, and 99bp when Exon 3 is lacking.

### Behavioral Testing

All mice were moved into a reverse light-cycle housing/testing room (lights off at 10 AM) at 5 weeks and allowed to acclimate for 2-3 weeks before the start of experiments. Experiments were performed in the housing room (ethanol drinking) or an adjoining room [loss of the righting response (LORR), rotarod]. Mice were group-housed 4 to 5 per cage based on genotype and sex. The same mice were sequentially tested on the rotarod, LORR, and drinking assays, with 4-7 days between assays.

### Drugs

Injectable ethanol solutions were prepared fresh daily in 0.9% saline (20%, v/v). Ethanol (Decon Laboratories, Inc.) was injected intraperitoneally (i.p.) at 0.02mL/g of body weight.

### Rotarod

In order to assess ethanol-induced motor ataxia, mice were trained on a fixed speed rotarod (Ugo Basile, Gemonio, Province of Varese, Italy) at 11rpm. Training was considered complete when mice were able to remain on the rotarod for 60sec. Following training, mice were injected with ethanol (2g/kg, i.p.) and every 15min mice were placed back on the rotarod and latency to fall was measured until mice were able to remain on the rotarod for 60sec.

### Loss of the Righting Response (LORR)

Sensitivity to the sedative/hypnotic effects of ethanol was determined using the LORR assay. Mice were injected with ethanol (3.5g/kg, i.p.) and when mice became ataxic, they were placed in the supine position in V-shaped plastic troughs until they were able to right themselves 3 times within 30sec. LORR was defined as the time from being placed in the supine position until they regained their righting reflex. Body temperatures were maintained using a heat lamp throughout the assay.

### Two-Bottle-Choice Every-Other-Day (2BC-EOD) Drinking

Mice were given access to ethanol (15%, v/v) and water for 24hr sessions every other day for 12 days starting at 12 PM. Water alone was offered on off days. Purchased drinking bottles were 15mL with 3.5-inch sipper tubes (Amuza, San Diego). The side placement of the ethanol bottles was switched with each drinking session to avoid side preference. Ethanol solutions were prepared fresh daily. Bottles were weighed before placement and after removal from the experimental cages. Empty cages with sipper bottles were used to control for leakage, and leakage amount was subtracted from amount consumed by the mice. The quantity of ethanol consumed, and total fluid intake, was calculated as g/kg body weight per 24hr. Preference was calculated as amount ethanol consumed divided by total fluid consumed per 24hr.

### Statistical Analysis

Statistical analysis was performed with GraphPad Prism (GraphPad Software, Inc., La Jolla, CA) for two-tailed Mann-Whitney test and two-way ANOVA (with mixed-effects analysis (i.e. when technical failures are present), multiple comparisons, and repeated measures when appropriate). Significant main effects were subsequently analyzed with Benjamini, Krieger, and Yekutieli two-stage linear step up procedure post-hoc analysis^10^. Technical failures were approprietely removed from analysis.

The two control groups were first compared to one another; if no difference was found between control groups, these groups were pooled and tested against the MyD88 KO group. Graphs show control groups plotted separately even though they were analyzed together, unless noted otherwise. Because of well-known sex differences on the behaviors of interest, and because male and female mice were tested on separate days, each sex was analyzed separately. Statistical significance was defined as p ≤ 0.05 and q ≤ 0.05. All data are presented as mean ± S.E.M.

## Results

### CRISPR/Cas9-mediated Mutagenesis

Preliminary testing of the CRISPy TAKO method occurred *in vitro* using embryos electroporated at the one-cell stage, cultured until the blastocyst stage, then genotyped (Fig. 2). To enhance CRISPR mutagenesis frequency, each gene was targeted simultaneously with four gRNAs that were tiled across a small section of the gene. In addition, we tested two concentrations of Cas9 protein (100 and 200ng/µL) that were higher than the minimum amount we typically use (i.e., 50ng/µL).

The first gene targeted was an unannotated ethanol-responsive gene, *4930425L21Rik*, using 4 gRNAs that span ∼400bp of the putative promoter and first Exon (Fig. 2A). Agarose gel electrophoresis of PCR amplicons that span the targeted locus indicated that 3 of 5 embryos tested at 100ng/µL Cas9 had obvious indels whereas 2 embryos (#’s 1.2 and 1.3; Fig. 2A, B) had amplicons that were grossly indistinguishable from the 613bp WT control amplicon (Fig. 2B). Sanger sequencing revealed #1.2 as heterozygous for WT and a 21bp deletion (Fig. 2A). At 200ng/µL Cas9 protein, all seven embryos assessed were found to harbor deletions of varying sizes (Fig. 2A, B). Thus, 0% of the embryos electroporated with 200ng/µL Cas9 harbored WT amplicons that were visible on the agarose gel or detectable by amplicon bulk sequencing.

The second gene targeted was another unannotated ethanol-responsive gene, *Gm41261*. Four gRNAs spanning ∼350bp within the putative first exon were used (Fig. 2C). Agarose gel electrophoresis of PCR amplicons that span the targeted locus indicated that 1 of 5 embryos tested at 100ng/µL Cas9 had an obvious indel (#5.6; Fig. 2C, D), whereas the other 4 of 5 embryos had amplicons that were indistinguishable from the 506bp WT control amplicon (Fig. 2D). Sanger sequencing revealed one embryo (#5.3; Fig. 2C, D) was homozygous WT, whereas the other four embryos harbored various small deletions (Fig. 2C). At 200ng/µL Cas9, all six embryos assessed were found to harbor deletions of varying sizes (Fig. 2C, D). Although one embryo (#7.5) had a PCR product approximately the size of the WT amplicon (506bp), Sanger sequencing revealed a 14bp deletion. Sanger sequencing also revealed a sequence inversion in #8.1, along with a 16bp insertion directly following the inverted sequence (Fig. 2C). Thus, 5 of 6 embryos electroporated at 200ng/µL Cas9 protein did not harbor detectable WT amplicons by agarose gel or amplicon bulk sequencing. Because the higher 200ng/µL Cas9 concentration showed greater mutagenic activity in both *4930425L21Rik* and *Gm41261*, this concentration was utilized in targeting MyD88.

As proof-of-concept and to validate our method *in vivo*, we created MyD88 CRISPy TAKO mice. The four gRNAs were tiled across a 209bp region of *MyD88* that included Exon 3 (Fig. 3A). Exon 3 was targeted because prior traditional global MyD88 KO studies demonstrated that deletion of this Exon inactivates MyD88^11^ and imparts an alcohol behavioral phenotype^5-7^.

Implantation of embryos electroporated with *MyD88* gRNAs yielded 54 offspring (n = 26 females, n = 28 males). Thirty-one offspring (n = 16 females, n = 15 males) were derived from electroporation of Mock-treated control embryos that were handled identically except that the crRNAs were omitted from the electroporation solution. All mice born from electroporated embryos were genotyped for gross indels at *MyD88* Exon 3 using endpoint PCR. The 494bp WT PCR amplicon was invariant and readily detectable in Jax and Mock-treated control samples as expected (Fig. 3B and data not shown). In stark contrast, 52 of 54 MyD88 KO mice displayed gross indels that were readily apparent following gel electrophoresis of PCR products. PCR results are shown in Figure 3B for the subset (n = 15/sex) of the MyD88 mutant mice created that were selected for behavioral phenotyping. The indels varied from animal to animal and were approximately 50-300bp smaller than the 494bp WT amplicon. To accurately characterize the mutations present, we sequenced the PCR products of the mutated mice selected for behavior As illustrated in Figure 3A, deletions removed Exon 3, deleted spice junctions, and/or are predicted to create frameshifts.

Cerebellar tissue from a random subset of MyD88 CRISPy TAKOs and controls were used for RT-PCR analysis using PCR primers that bind to Exons 2 and 4 to examine MyD88 mRNA. This analysis revealed the expected 280bp fragment in Jax and Mock WT control samples (Fig. 3C). In contrast, none of the six MyD88 CRISPy TAKO mice examined peoduced a fragment of this size. Five of the six samples produced a predominant band of ∼99bp was would be expected for MyD88 mRNA that lacked Exon 3. One sample produced a major band of ∼210bp and may represent a splicing defect. Thus, MyD88 CRISPy TAKO mice are likely to be functional KOs.

### Ethanol-Induced Loss of Righting Response (LORR)

No difference in ethanol LORR (3.5g/kg, i.p.) was found between the Mock and Jax control groups for males or females (p = 0.9671 and p = 0.7345, respectfully; Fig. 4). Therefore, Mock controls and Jax controls were combined and compared to MyD88 KOs (for completeness, control results are plotted separately). Male MyD88 KOs exhibited a significant reduction in ethanol-induced LORR duration when compared to controls (p = 0.0068; Fig. 4A). No difference was observed in females for ethanol-induced LORR (Fig. 4B).

**Figure 4:**
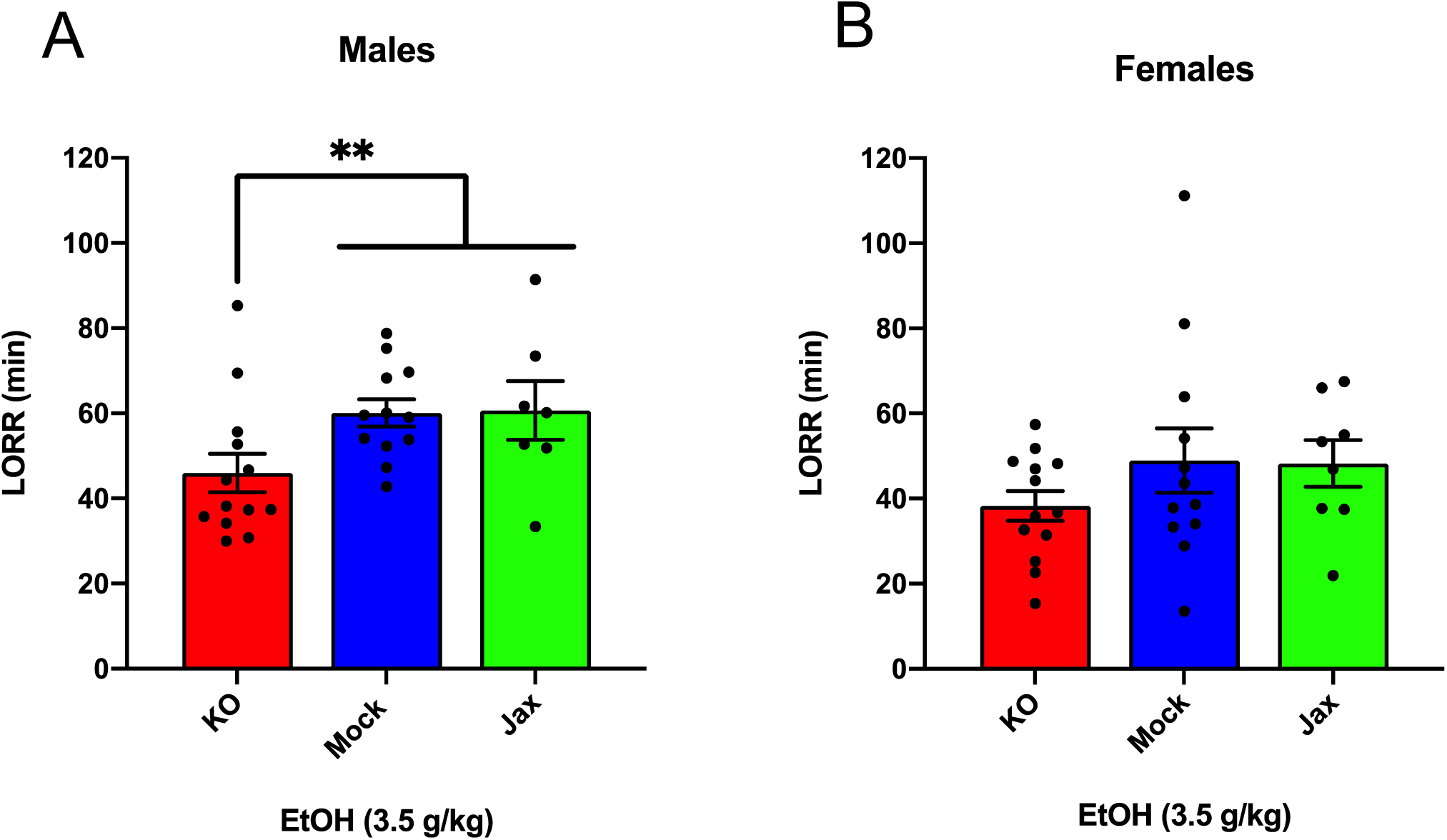
Duration of LORR induced by ethanol (3.5 g/kg, i.p.) injection in male **(A)** and female **(B)** MyD88 KOs, Mock controls and Jax controls. For both males and females, mock controls and Jax controls did not differ and therefore were pooled and compared to same sex MyD88 KOs (data are plotted separately for completeness). KO males had reduced duration of LORR compared to the combined control group (**p < 0.01). For females, no difference was observed between MyD88 KOs and the combined control group. Values represent mean ± SEM. Data were analyzed using a two-tailed Mann-Whitney test.

### Ethanol-Induced Motor Incoordination

The ataxic effects of an acute ethanol injection (2g/kg, i.p.) were measured using a constant speed (11rpm) rotarod test. For male mice, comparison of Mock and Jax controls showed a significant effect of time [F (2.281, 41.05) = 36.41, p < 0.0001], and time x genotype [F (9, 162) = 3.209, p = 0.0013], but no effect of genotype (Fig. 5A). Because of the time x genotype interaction, control groups were not combined for this analysis. Repeated measures two-way ANOVA of all three groups revealed a significant effect of time [F (2.474, 71.73) = 59.01, p < 0.0001], and an effect of time x genotype [F (18, 261) = 1.964, p = 0.0120] but no effect of genotype (Fig. 5A). Posthoc comparisons revealed that male Mock control mice recovered more quickly than Jax controls at the 15min timpoint (q = 0.0042).

**Figure 5:**
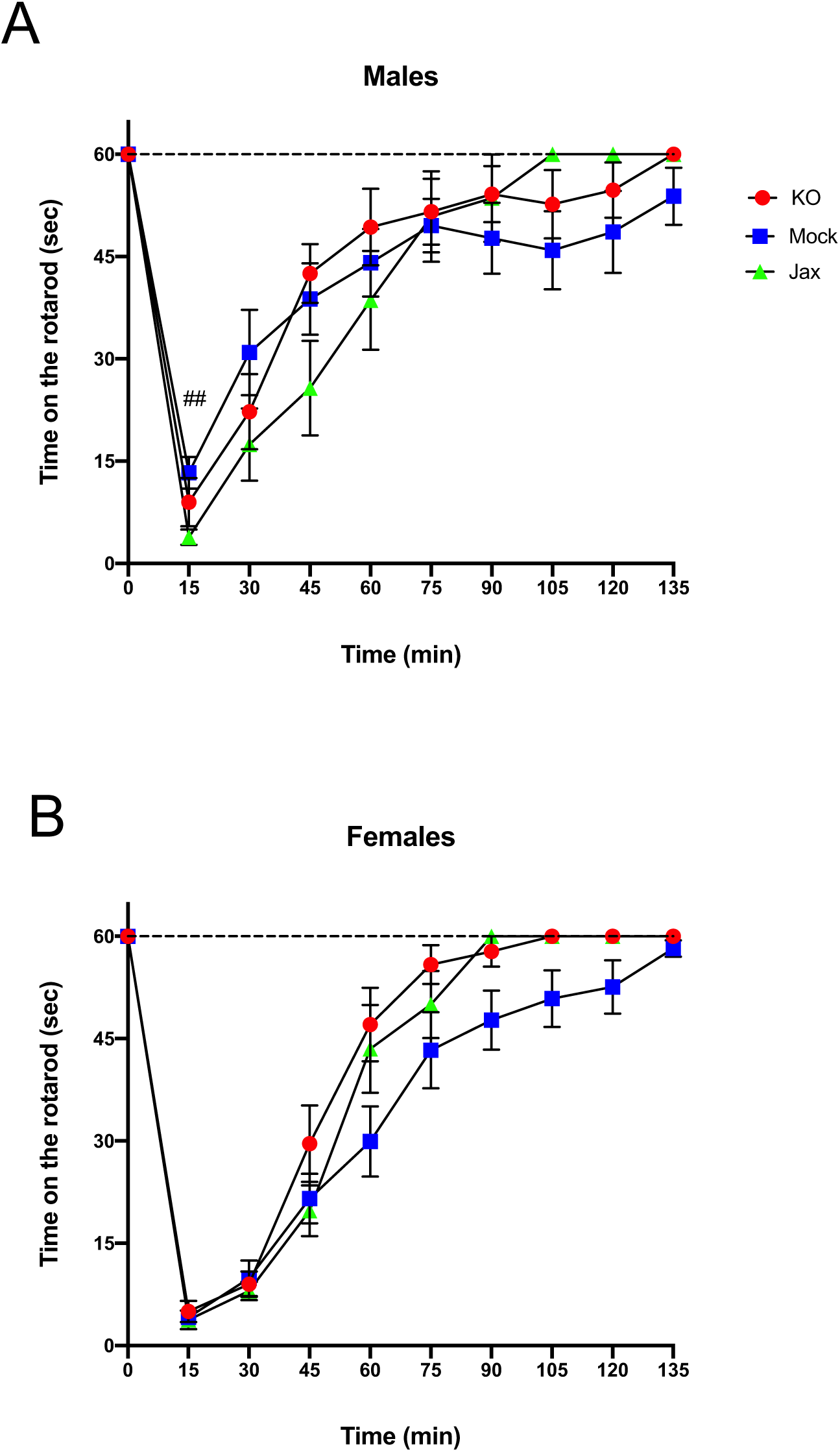
Recovery from ethanol (EtOH)-induced motor incoordination in MyD88 KO and control mice. **(A)** A significant time x genotype interaction was observed for male mock controls and Jax controls. Therefore. control groups were not combined. Jax controls recovered more slowly compared to Mock controls at the 30 min timepoint (##q < 0.01). **(B)** Female mock controls and Jax controls did not differ and were combined for comparison to MyD88 KOs (data are plotted separately for completeness). Post-hoc analysis did not reveal any significant differences between groups at any time points. Data represent time in seconds on the rotarod after injection of EtOH (2 g/kg, i.p.). Values represent mean ± SEM. Data were analyzed by repeated-measures 2-way ANOVA with multiple comparisons followed by Benjamini, Krieger, and Yekutieli post hoc-test.

For females, comparison of Mock and Jax controls revealed a significant effect of time [F (2.775, 55.51) = 89.05, p < 0.0001], but no effect of genotype or time x genotype (Fig. 5B). Therefore, the two control groups were combined (data plotted separately for completeness) and compared to Myd88 KOs. There was a significant effect of time [F (2.664, 87.90) = 148.3, p < 0.0001], and genotype [F (1, 33) = 4.721, p = 0.0371], but no effect of the time x genotype interaction (Fig. 5B). Post-hoc analysis did not reveal any significant differences.

### Two-Bottle Choice Every-Other-Day (2BC-EOD) Drinking

Mice were tested for ethanol drinking using an intermittent every-other-day, two bottle free choice consumption assay over a period of 12 days. Mock and Jax control groups were first compared against each other; two-way ANOVA mixed-effects analysis was used for all 2BC-EOD statistical analyses. For males, there was a significant effect of time for ethanol intake [F (3.333, 62.00) = 5.740, p = 0.0011], ethanol preference (2.702, 50.27) = 10.85, p < 0.0001], and total fluid intake [F (3.392, 63.09) = 17.98, p < 0.0001], but there was no effect of genotype or time x genotype interaction for any of these parameters (Fig. 6A-C). Therefore, both Mock and Jax control groups were combined (data plotted separately for completeness) and compared to MyD88 KOs.

**Figure 6:**
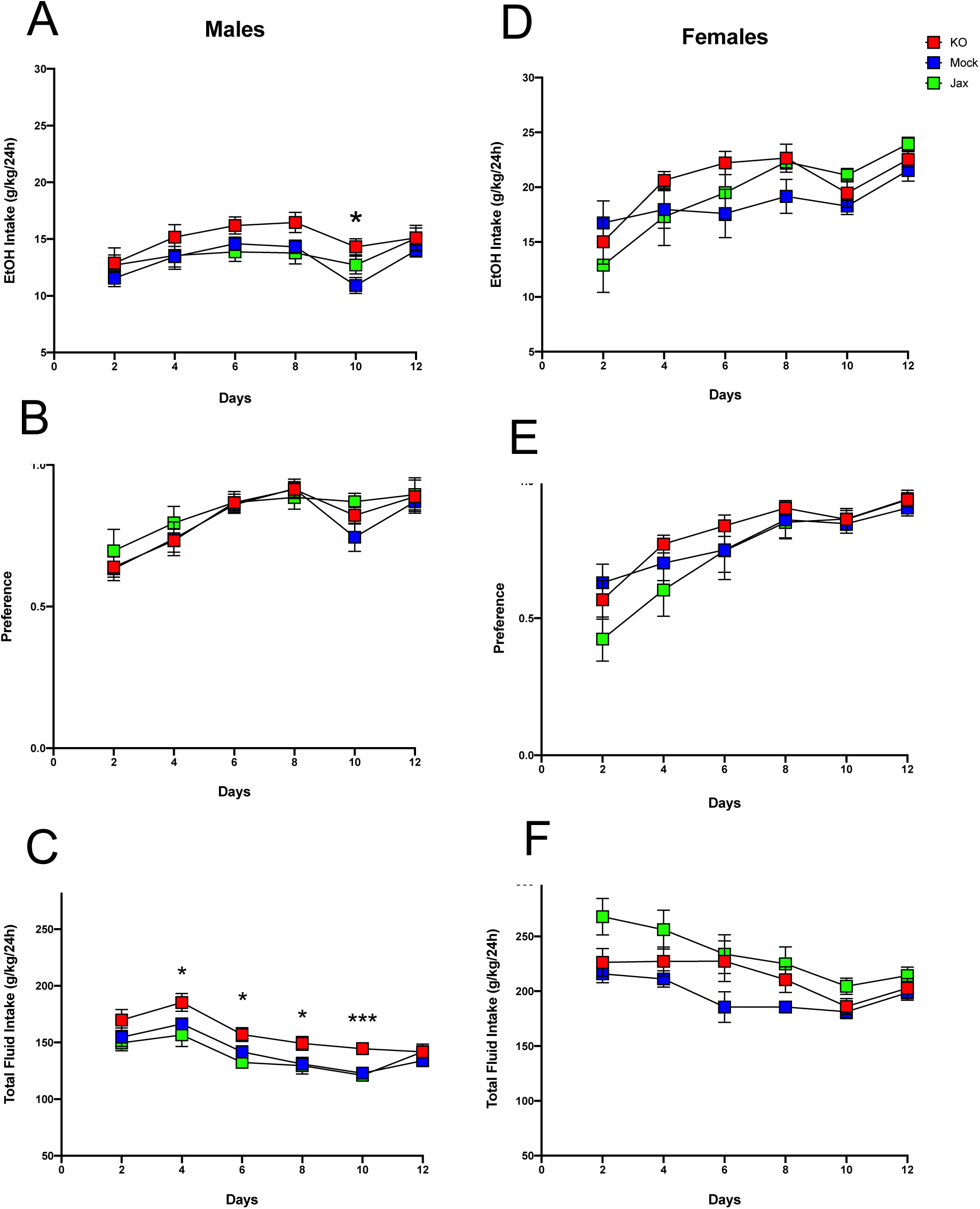
Two-bottle choice, every-other-day drinking in MyD88 KO, Mock control and Jax control mice. Left, males; right, females. **A and D**. Ethanol (EtOH) intake (g/kg/24 h), **B and E**. EtOH preference. **C and F**. Total fluid intake (g/kg/24 h) in control (Jax C57BL/6J, n = 8; Mock-treatment control, n = 13-14) versus mutant mice (n = 13-14 MyD88 KO). Values represent mean ± SEM. Data were analyzed by repeated-measures 2-way ANOVA (mixed-effects analysis where appropriate) with multiple comparisons followed by Benjamini, Krieger, and Yekutieli post-hoc tests (*p < 0.05 and ***p < 0.001 between MyD88 KO and combined controls.

Analysis of ethanol intake in males between the combined control group and MyD88 KO revealed a main effect of time [F (4.123, 129.5) = 10.67, p < 0.0001] and genotype [F (1, 32) = 4.850; p = 0.0350], but no interaction between the two (Fig. 6A). Post-hoc analysis revealed that MyD88 KO males had significantly greater intake on day 10 (q = 0.0285) compared to controls. For preference in males, there was an effect of time [F (3.365, 105.7) = 24.02, p < 0.0001] but no effect of genotype or time x genotype (Fig. 6B). For total fluid intake in males, there was a main effect of time [F (3.915, 122.9) = 36.79, p < 0.0001] and genotype [F (1, 32) = 8.897, p = 0.0054], but no time x genotype interaction between MyD88 KO and controls (Fig. 6C). Post-hoc analysis revealed significantly increased total fluid consumption in MyD88 KOs vs controls on days 4, 6, 8 (q = 0.0118), and day 10 (q = 0.0002) (Fig. 6C).

In females, Mock and Jax control groups were first compared. There was a significant effect of time on ethanol intake [F (2.412, 47.27) = 8.979, p = 0.0002] and ethanol preference [F (2.626, 51.47) = 22.58, p < 0.0001], but no effect of genotype or time x genotype interaction on either parameter (Fig. 6D, E). Therefore, both control groups were combined (data plotted separately for completeness) and compared to MyD88 KOs. For ethanol intake, females showed a main effect of time [F (2.632, 86.85) = 12.50, p < 0.0001] but not genotype or time x genotype interaction (Fig. 6D). Similarly, for ethanol preference a significant effect of time [F (3.317, 109.5) = 29.10, p <0.0001] but not genotype and time x genotype interaction was observed (Fig. 6E). Thus, ethanol intake and preference did not differ between female MyD88 KOs and the combined control group.

Comparing total fluid intake in female Mock and Jax controls revealed significant main effects of time [F (2.818, 55.22) = 9.800, p < 0.0001] and genotype [F (1, 20) = 10.41, p = 0.0042], but no time x genotype interaction (Fig. 6F). Therefore, for female total fluid intake, control groups were not combined and each genotype was considered separately. This analysis revealed a significant effect of time [F (2.650, 84.79) = 11.20, p < 0.0001] and genotype [F (2, 33) = 4.221, p = 0.0223] (Fig. 6F), but no time x genotype interaction. Post-hoc analysis did not reveal any significant differences.

## Discussion

The current study reports on a CRISPR/Cas9-mediated mutagenesis protocol that is suitable for rapid screening for the phenotypic effects of gene KO *in vivo*. Traditional CRISPR mouse KO procedures require three generations of animal breeding and genotyping, which is time consuming and expensive. In contrast, with the CRISPy TAKO protocol described here, first generation gene-targeted F0 mice can be rapidly produced and screened for phenotypic effects. Although individual F0 mice harbor a variety of mutant alleles for the gene of interest, careful project design ensures that each F0 animal is functionally equivalent, i.e., a gene KO. F0 animals can be directly screened for phenotypes of interest. If no phenotype is detected, the gene is rapidly eliminated from further consideration. If an interesting phenotype is observed, a F0 animal can be bred as in traditional approaches to establish a true breeding line that can first be tested to confirm the phenotype. This will ensure rigor and reproducibility in the experimental pipeline. Subsequently, the line can be maintained long-term and more detailed, rigorous mechanistic studies can be conducted. The CRISPy TAKO approach can save valuable time and minimize animal numbers and financial resources.

There are several keys to the success of this approach. First, we use embryo electroporation to facilitate genetic modification of a large number of animals with minimal effort. Large numbers of embryos (n = 30-50) can be simultaneously transfected with CRISPR reagents at a very high efficiency^12-16^. This avoids the limiting bottleneck of directly injecting each individual embryo. Second, achieving a very high level of indel formation that ablates function of the gene of interest in each animal is critical. We observed that 52 of 54 of animals harbored inactivating indels, while the other two harbored small mutations that were not charecterized. To achieve this high KO efficiency, we simultaneously utilized four gRNAs that targeted a small, functionally important portion of the gene of interest. In other experiments, we have observed that the mutagenesis efficiency of a single gRNA is highly variable. Simultaneous use of two gRNAs tends to increase mutagenesis efficiency. We reasoned that an even higher number of gRNAs tiled across a small but functionally important part of a gene would result in even higher efficiency. We are unsure if four is the optimal number of gRNAs, but this should be rigorously explored in future studies. For *in vivo* proof-of-concept, we focused on a small, single Exon of the *MyD88* gene that results in a null allele when disrupted^5-7,11^. This approach should also work by targeting the promoter or any region that is critical for function of the gene and/or gene product. It should also be noted that we observed that 200ng/µL Cas9 protein in the electroporation mix produced a much higher rate of indel formation compared to 100ng/µL. While 200 ng/µL is 4x the minimum amount we typically use in our lab for most CRISPR embryo electroporation experiments, this amount is less than that reported in the literature^15,16^.

The CRISPy TAKO approach could be further streamlined and throughput increased if control animals for comparison could be procured directly from a vendor. However, it is conceivable that the *in vitro* embryo manipulation / CRISPR electroporation procedure could introduce some unknown variable that could impact the phenotype of interest regardless of the gene targeted for modification. Therefore, we compared phenotypes of control animals procured directly from JAX with isogenic controls that were produced in-house, in parallel to the MyD88 TAKOs. This in-house control group was created using procedures that were identical to those used to create MyD88 TAKOs except that crRNAs were omitted from the electroporation reactions. We observed near complete concordance between these Mock controls and Jax control animals for the behavioral phenotypes of interest. We only observed a subtle female-specific difference in total fluid consumption in the 2BC-EOD assay (Fig. 6G) and a male-specific genotype x time interaction on the rotarod (Fig 5A). We conclude that mice purchased directly from a vendor can be used as a control group for screening CRISPy TAKO mice for behavioral alterations provided the controls are the same genetic background as those animals that served as embryo donors. Using a single vendor-derived control group will substantially increase throughput and reduce expenses.

As proof-of-concept of the CRISPy TAKO approach, we focused on *MyD88* as a candidate gene. We sought to functionally validate our approach by comparing behavioral phenotypes observed with those previously reported for global MyD88 KO mice produced using traditional gene targeting technology, which displayed robust alterations in ethanol-induced behavioral responses and ethanol drinking behavior^5-7^. Overall, similar behavioral results were observed between traditional MyD88 KOs and MyD88 CRISPy TAKOs (Table 1). Consistent with previous findings, the MyD88 KO females show faster recovery time from ethanol’s incoordination effects (Fig. 5B), but contrary to those studies, no difference between male MyD88 KOs and controls is reported here (Fig. 5A). Also consistent with previous reports, albeit with a milder effect size^5^, MyD88 KO males had greater consumption of ethanol than controls (Fig. 6A). However, KO males in the present study did not have a difference in preference when compared to controls (Fig. 6B), but had significantly increased total fluid intake compared to controls (Fig. 6C), suggesting these male mice drink more fluid in general, and it is not specific to ethanol. Altered total fluid intake in MyD88 KO females compared to controls (Fig. 6F) is consistent with the published literature^5^.

It is unclear if the results presented here show a milder phenotype than those previously reported^6,7^, or if these differences are simply due to experimental variation that is common in behavioral studies between labs, facilities, and universities^17,18^. Although all studies were conducted using C57BL/6J mice, the current study utilized mice sourced directly from JAX, whereas Blednov et al., 2017a/b used mice sourced from JAX that were bred in-house for an unspecified number of generations. It is also possible that the CRISPy TAKO approach is slightly less sensitive than the traditional KO approach for detecting phenotypic changes. The most likely explanation for such a possibility is that F0 CRISPR mice may be mosaic and it is conceivable that tail DNA genotyping is not reflective of the genetic changes that occur in the brain of the mutant animals. It is possible that some WT MyD88 is expressed in the brain of some F0 animals, however this is unlikely because RT-PCR analysis of a subset of TAKO MyD88 mice did not reveal WT bands (Fig. 3C). Nonetheless, the approach described here should be very useful for a first pass screening method to identify genes with a large effect on a phenotype of interest. The usefulness of this approach for detecting subtle genotypic differences requires further evaluation.

One limitation of the approach as outlined is the potential for off-target effects of CRISPR mutagenesis. This approach uses multiple gRNAs simultaneously along with a relatively high concentration of Cas9, both of which could lead to off-target effects. Although off-target effects were not examined in this study, they are unlikely to explain the phenotypic changes we observed. The main behavioral phenotypes observed in the MyD88 CRISPy TAKO mice are the same as those observed in MyD88 global KOs that were produced using traditional, non-CRISPR gene targeting techniques. Furthermore, several studies have reported that off-target effects in CRISPR/Cas9 animals is minimal with careful selection of gRNAs as done in the present study^19-21^.

In summary, we propose using the CRISPy TAKO approach for rapidly screening large numbers of genes *in vivo* to identify those that have large effects on a phenotype of interest. Once such a gene is identified, an individual animal that harbors a confirmed KO allele should be mated to establish a true breeding mutant KO line. A true breeding line will be useful for future studies to (a) confirm the phenotype of interest, (b) to test for and rule out the potential impact of off-target mutations, (c) to enable the rigorous testing of control and KO littermates derived from heterozygous matings, and (d) to provide an unlimited source of uniform animals for further, in-depth analyses and long term line maintenance.

We conclude that the CRISPy TAKO method can be used for efficient, moderate throughput, *in vivo* screens to identify genes that impact whole animal responses when ablated. This method avoids the extensive animal breeding, time, and resources required with traditional CRISPR animal KO approaches. This method should find widespread use in studies where moderate to large numbers of genes must be rapidly screened for effects that cannot be interrogated *in vitro*, such as whole animal behavioral responses.

## Author Contributions

Project conception and gRNA design divised by GEH. *In vitro* analysis conducted by AS and GEH. *In vivo* project design, organization, and analysis conducted by SLP. SLP and AS managed the behavioral experimentation together. All authors contributed to writing and editing of the manuscript.

## Acknowledgements

The authors would like to acknowledge Carolyn Ferguson for expert technical support, Tanya Kenkre, PhD for statistical consultation, and members of the INIA-Neuroimmune consortium for helpful discussions and constant encouragement.

