## Supplemental Table 1 for "CRISPR Turbo Accelerated Knock Out (CRISPy TAKO) for rapid *in vivo* screening of gene function"

| Name | Sequence |
|---|---|
| *4930425L21Rik* #1 gRNA | AGACACTAATATTGCAGACG AGG |
| *4930425L21Rik* #2 gRNA | TTATTTTTCTGCAAGGGGTT GGG |
| *4930425L21Rik* #3 gRNA | ACACTATCGACCTAATAGCT AGG |
| *4930425L21Rik* #4 gRNA | ATTTTAAACCCTCTGTTACT TGG |
| *Gm41261* #1 gRNA | CAATTTGCAATTCTCTTCCA GGG |
| *Gm41261* #2 gRNA | AGAATAAACAGGTGTGACGG TGG |
| *Gm41261* #3 gRNA | CTTATCAGGTCTTTGATCAG AGG |
| *Gm41261* #4 gRNA | GGTCTTTTACTTTTCTCTTT AGG |
| MyD88 T3 gRNA | CCTTTTCTCAATTAGCTCGC TGG |
| MyD88 T5 gRNA | GCACAAACTCGATATCGTTG GGG |
| MyD88 T15 gRNA | AGGTTGGTTAAACATCTAAG AGG |
| MyD88 T30 gRNA | GGCGTTTGTCCTGAGGACAG GGG |
| *4930425L21Rik* F5 PCR primer | GTGTCCAGCATTGTGCCAAG |
| *4930425L21Rik* R5 PCR primer | TCTAAAAGGGGCCCTCCAGT |
| *Gm41261* F10 PCR primer | CTCACCAAAATTCAACCTGGAG |
| *Gm41261* R10 PCR primer | GCTTCAGAGCTCACTGGTGT |
| MyD88 F1 PCR primer | CCGGGATTTCATCTGGGAGG |
| MyD88 R1 PCR primer | ACTGCGGTGACTTCCTTCAG |
| MyD88 F2 PCR primer | GGTGGCCAGAGTGGAAAGCAGTGTCCC |
| MyD88 R2 PCR primer | GAAACAACCACCACCATGCGGCGACA |

Supplemental Table 1. gRNA target sites and PCR primer sequences. All sequences are written in a 5’ to 3’ orientation. Note: underlined sequence in each gRNA target site is the protospacer adjacent motif.
